# Serotonin 5-HT_2C_ receptor as a cellular target of PI3K inhibitor LY294002 and its analog LY303511

**DOI:** 10.1101/2025.11.06.687025

**Authors:** Polina D. Kotova, Ekaterina A. Dymova, Oleg O. Lyamin, Olga A. Rogachevskaja, Evgeniya A. Voronova, Stanislav S. Kolesnikov

## Abstract

The inhibitory analysis of intracellular signaling pathways is widely employed to gain insight into molecular mechanisms underlying diverse physiological processes. Unfortunately, the essential drawback of this basically effective methodology is that many, if not all, inhibitors, antagonists, modulators, and blockers can affect cellular functions not only acting through specified cellular targets, but also causing off-target effects. In particular, the class I phosphatidylinositol-3-kinase (PI3K) inhibitor LY294002 and its PI3K-inactive structural analog LY303511 have been shown to affect agonist-induced Ca^2+^ signaling in cells of various types independently of PI3K activity. Here we studied serotonin-induced Ca^2+^ signaling in HEK293 cells expressing the recombinant mouse 5-HT_2C_ receptor and analyzed the effects of LY294002 and LY303511 on cell responsiveness. As shown with Ca^2+^ imaging, both LY294002 and LY303511 affected intracellular Ca^2+^ but via distinct mechanisms. LY294002 suppressed responsiveness of assayed cells to serotonin in a manner suggesting that this substance acted as a competitive antagonist of the 5-HT_2C_ receptor. In turn, LY303511 itself triggered Ca^2+^ transients in 5-HT_2C_-positive cells, exhibiting traits of a 5-HT_2C_ agonist. In support of these findings, molecular docking and molecular dynamics simulations validated the binding of both LY294002 and LY303511 to the 5-HT_2C_ receptor and occupying its orthosteric site. Altogether, physiological findings and computational data suggested that the observed effects of these compounds were most likely mediated by extracellular mechanisms associated with the direct interaction of both with the 5-HT_2C_ receptor. This expands the list of non-specified cellular targets of LY294002 and LY303511 with 5-HT_2C_ subtype of serotonin receptors.

## 1. Introduction

Intracellular signaling relies on precise molecular interactions which are commonly transient and spatially compartmentalized, and also can depend on crosstalk between several pathways. Therefore, signaling and regulatory processes could not be completely revealed and precisely characterized beyond the intracellular context. Being combined with the inhibitory analysis, the direct monitoring of physiological processes in cells with electrophysiological or imaging methods is a rather effective approach for disclosing signaling pathways and evaluating contribution of particular regulatory molecules to their functionality. The inhibitory analysis is widely employed as providing transparent experimental logic and due to availability of numerous agonists, antagonists, activators, modulators and inhibitors targeting a huge number of different cell surface receptors and intracellular molecules.

Meanwhile, the essential drawback of this methodology is that many, if not all, compounds available for the inhibitory analysis can affect intracellular processes not only acting through specified cellular targets, but also causing off-target effects. For instance, the phosphodiesterase inhibitor IBMX also serves as an adenosine receptor antagonist.^1^ The mitogen-activated protein kinase kinase (MAPKK) inhibitor PD98059 has the antagonistic effect towards aryl hydrocarbon receptor.^2^ U73122 can affect intracellular Ca^2+^ signaling not only through the inhibition of its primary target phospholipase C (PLC) but also by modulating certain Ca^2+^ permeable channels.^3,4^ The ubiquitously used activator of adenylate cyclase forskolin directly inhibits certain voltage-gated K^+^ channels.^5^ Dihydropyridine, a Ca^2+^ channel blocker, stimulates the Ca^2+^-sensing receptor.^6^ Reportedly, inhibitors of class I phosphatidylinositol-3-kinase (hereafter referred to as PI3K) also can exert off-target effects.^7–10^

The growing body of evidence points at PI3Ks as a master regulators of a variety of physiological processes such as growth, cell survival and migration, and intracellular vesicular transport.^11^ The PI3K family comprises three classes of intracellular lipid kinases, all of which phosphorylate phosphatidylinositides at the 3’ position of the inositol ring, but differ in substrate specificity and sequence homology.^12^ The PI3K phosphorylates phosphatidylinositol (3,4)-bisphosphate (PIP_2_) at the plasmalemma, producing phosphatidylinositol (3,4,5)-trisphosphate (PIP_3_) and modulating activity of PIP_3_-dependent enzymes and related signaling pathways. Notably, the PI3K/Akt pathway has been shown to regulate Ca^2+^ signaling.^13–17^ The role of PI3K in intracellular processes has largely been verified with PI3K inhibitors, including wortmannin and also LY294002 (LY294) alone or in combination with LY303511 (LY303), a structural analog of LY294 inactive against PI3K.^18,19^ As reported later, however, LY294 and its analogs could affect agonist-induced Ca^2+^ signaling not necessarily via the PI3K inhibition but involving unrelated mechanisms. It was particularly shown that LY294 and its analogs suppressed Ca^2+^ signals initiated by acetylcholine, histamine, and serotonin in assayed cells independently of the PI3K inhibition.^7–10^ Although in the mentioned studies, veritable mechanisms mediating the LY294 effects were not established with confidence, some findings pointed to appropriate aminergic G-protein-coupled receptors (GPCRs) to be likely cellular targets of the PI3K inhibitors.^9, 10^

Here we studied Ca^2+^ signaling induced by serotonin in the HEK293 cells expressing the recombinant mouse serotonin 5-HT_2C_ receptor and analyzed the effects of LY294 and LY303 on cell responsiveness. As demonstrated with Ca^2+^ imaging, LY294 did not affect Ca^2+^ in resting cells but suppressed cell responsiveness to serotonin. This effect was not mediated by inhibition of either PI3K or key molecules of phosphoinositide signaling cascade (Gq, PLC and IP_3_Rs), but it was attributed to a direct action of LY294 on the 5-HT_2C_ receptor as a competitive antagonist. In contrast, LY303 itself induced Ca^2+^ transients in 5-HT_2C_-positive cells, but not in non-transfected cells, exhibiting traits of a 5-HT_2C_ agonist. Molecular docking and molecular dynamics (MD) simulations pointed out that both LY294 and LY303 could be bound to the 5-HT_2C_ receptor in its orthosteric site. Thus, LY294 and LY303 appear to be ligands of the 5-HT_2C_ receptor, acting extracellularly as an antagonist and an agonist, respectively.

## 2. Materials and Methods

### 2.1 Cell lines and culturing

This study relied on HEK293 cells and their derivative HEK/5-HT_2C_ cells, that stably express the mouse serotonin 5-HT_2C_ receptor (see Supplemental methods 1 and Supplemental Fig. 1B–D for details of the HEK/5-HT_2C_ cells generation and characterization). Cells were cultured in the Dulbecco’s modified Eagle’s medium (DMEM; Servicebio) contained 10% (vol/vol) fetal bovine serum (HyClone), 1% glutamine, 0.1 mg/ml gentamicin (Gibco), and selective antibiotic G-418 (0.3 mg/ml) for HEK/5-HT_2C_ cells (Invivogen). Cells of all lines were grown in 12-well culture plates in a humidified atmosphere of 5% CO_2_ at 37°C.

### 2.2 Ca^2+^ imaging

Immediately before the experiments, cultured cells were washed with the Versene solution (Sigma-Aldrich), incubated in the 0.25% Trypsin-EDTA solution (Sigma-Aldrich) and resuspended in a complete growth medium. Isolated cells were plated onto a hand-made photometric chamber, the bottom of which was coated with Cell-Tak (Corning), ensuring sufficient cell adhesion. The chamber of nearly 150 μl volume was a disposable coverslip (Menzel-Glaser) with an attached ellipsoidal wall made of thermo-glue.

For Ca^2+^ imaging, attached cells were loaded with the Ca^2+^ dye Fura-2 by incubating them in a bath solution containing 4 μM Fura-2 AM (AAT Bioquest) and 0.02% Pluronic F-127 (SiChem) at room temperature (23 – 25 °C) for 20 min. Next, cells were rinsed several times, and kept in the bath solution for at least 1 h prior to recordings. The bath solution contained (mM): 130 NaCl, 5 KCl, 2 CaCl_2_, 1 MgCl_2_, 10 glucose, 10 HEPES (pH 7.4). When necessary, 2 mM CaCl_2_ in the bath was replaced with 0.5 mМ EGTA + 0.4 mМ CaCl_2_, thus reducing free Ca^2+^ to nearly 260 nM at 23 °С as calculated with the Maxchelator program (http://maxchelator.stanford.edu). The used salts and buffers were from Sigma-Aldrich.

For monitoring Ca^2+^ signals initiated by IP_3_ uncaging, attached cells were loaded with both the Ca^2+^ dye Fluo-8 and caged-Ins(145)P3, a photoconvertible IP_3_ precursor. Cells were incubated in the bath solution in the presence of 4 μM Fluo-8 AM (AAT Bioquest), 4 μM caged-Ins(145)P3/PM (SiChem) and 0.02% Pluronic F-127 (SiChem) at room temperature for 20 min, rinsed several times, and kept in the bath solution for at least 1 h prior to recordings.

Experiments were carried out using an inverted fluorescence microscope Axiovert 135 equipped with an objective Plan NeoFluar 20x/0.75 (Carl Zeiss), a digital EMCCD camera iXon 888 (Andor Technology), and a hand-made illuminator. The computer-controllable illuminator with a set of light-emitting diodes provided epi-illumination at multiple wavelengths through a bifurcational glass fiber. For Fura-2 visualization, two channels of the fiber transmitted excitation light at 340 ± 6 nm and 380 ± 6 nm, respectively, emission was collected at 510 ± 40 nm. Deviations of cytosolic Ca^2+^ in individual cells were presented as the ratio F_340_/F_380_, where F_340_ and F_380_ are the instant intensity of cell fluorescence exited at 340 and 380 nm, respectively. For simultaneous Fluo-8 visualization and caged-Ins(145)P3 photolysis, one channel of the fiber was connected to the illuminator, providing excitation of Fluo-8 at 480 ± 10 nm, while the dye emission was collected at 520 ± 20 nm. Another fiber channel was connected to a pulsed solid laser LCS-DTL-374QT (Laser-Export) generating UV light at 355 nm. This laser operated in a two-harmonic mode and generated not only UV flashes required for IP_3_ uncaging, but also emitted at 532 nm, the light penetrating into the Fluo-8 emission channel and resulting in optical artifacts during IP_3_ uncaging. Deviations of cytosolic Ca^2+^ in response to IP_3_ uncaging in individual cells were presented as the ratio ΔF/F_0_, where ΔF = F − F_0_, F is the instant intensity of cell fluorescence, F_0_ is the intensity of cell fluorescence obtained in the very beginning of a recording and averaged over a 20 s interval. Serial fluorescence images were captured every second and analyzed using imaging software NIS-Elements AR 5.30.01 (Nikon). All chemicals were applied by the complete replacement of the bath solution in a 150 μl photometric chamber for nearly 2 s using a perfusion system driven by gravity. For data and graphical analysis Sigma Plot 15 (Systat Software Inc) was used. N represents the number of cells from at least three experiments conducted on different days.

### 2.3 Drugs

LY294 (LY294002 hydrochloride) and LY303 (LY303511 hydrochloride) were from MedChemExpress; serotonin hydrochloride, insulin, m-3M3FBS, H-89 dihydrochloride, U73122, acetylcholine chloride, and RS-102221 hydrochloride were from Tocris Bioscience; wortmannin and ATP were from Sigma-Aldrich.

### 2.4 Quantifying of LY294 and LY303 effects

To characterize a dose dependence of LY294 effect on serotonin-induced Ca^2+^ transients, a fraction of cells responsive to 4 nM serotonin in the presence of LY294 at variable doses was evaluated. The IC_50_ value of LY294 was calculated by fitting this dose dependence using the Hill equation:

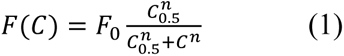

where F(C) is the fraction of cells responsive to 4 nM serotonin in the presence of LY294 at the given concentration C, F_0_ is the fraction of serotonin responsive cells in control, C_0.5_ is the LY294 concentration of the half maximal inhibitory effect (IC_50_), and n is the Hill coefficient.

To characterize the sensitivity of HEK/5-HT_2C_ cells to LY303, a fraction of cells generating Ca^2+^ transients to this compound at variable doses was evaluated. The EC_50_ value of LY303 was calculated by fitting this dose dependence using the equation:

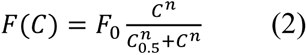

where F(C) is the fraction of cells responsive to LY303 at the given concentration C, F_0_ is the maximal fraction of LY303 responsive cells, C_0.5_ is the LY303 concentration of the half maximal effect (EC_50_), and n is the Hill coefficient.

### 2.5 Modeling of the mouse 5-HT_2C_ receptor structure

The 3D-structure of the mouse 5-HT_2C_ receptor is currently not available. Given that its amino acid sequence is 96% homologous to the human 5-HT_2C_ receptor, the crystal structure of the latter was used as a template for homology modeling of the mouse 5-HT_2C_ structure.

The human 5-HT_2C_ receptor structure, PDB ID: 6BQH,^20^ obtained from RCSB crystallographic database https://www.rcsb.org^21^ was modified by removing manually all non-protein molecules, e.g. the inverse agonist ritanserin and also a protein segment between the 5th and 6th transmembrane helices, which was added to facilitate the crystallization. The resultant gap at the intracellular loop 3 was filled in with the amino acid linker GSGSGSGS using the SWISS-MODEL tool.^22^ This sequence of glycine and serine residues was chosen due to their common occurrence in natural linkers, electroneutrality, and short side chains. In addition, similar sequences were previously used for modeling ligand-receptor interactions.^22^ The number of four GS-repeats was chosen to provide the necessary minimum that allowed for unconstrained dynamics of the 5th and the 6th helices of the human 5-HT_2C_ receptor.

Finally, the homologous modeling of the mouse 5-HT_2C_ receptor structure was accomplished using the SWISS-MODEL tool, based on the human 5-HT_2C_ receptor template described above.

### 2.6 Molecular docking

3D-structures of serotonin, wortmannin, hematein, LY294, and LY303 were obtained from the databases ZINC^23^ and Pubchem.^24^ All these structures were prepared for docking with AutoDockTools-1.5.6. Docking was performed with Autodock Vina 1.1.2.^25^ This docking tool allows constraining the search space of conformations energetically most favorable for forming a stable ligand-receptor complex by specifying an appropriate region on the receptor surface. The region between 5-HT_2C_ receptor transmembrane helices bounded by their extracellular loops and the receptor center looks reasonably adequate, given a number of experimentally-obtained crystal structures of serotonin receptors in complexes with their known ligands (e.g. with ritanserin, PDB ID: 6BQH).

Since no 3D-structure of the 5-HT_2C_ receptor in complex with serotonin is available now, the location of serotonin on the receptor was modeled similarly with docking and was then considered as the orthosteric serotonin-binding site.

### 2.7 Molecular dynamics

To eliminate steric clashes potentially produced by the docking, refine ligand pose on the receptor surface, and validate stability of the docking-predicted complexes, we performed molecular dynamics simulations using the pmemd.cuda tool, which is included in the Amber Molecular Dynamics Package (version 20),^26^ GPUs Nvidia 2080 and Nvidia 3080.

Protonation states of the 5-HT_2C_ receptor amino acid residues were preliminarily determined by PROPKA at pH 7.0,^27^ except for Asp100 which was protonated. The conserved Asp100 corresponds to Asp83 in rhodopsin, which is known to be protonated during the entire photocycle.^28^ Amber preparation tools charge terminal residues by default. To avoid that, N- and C-termini were capped with neutral acetyl and N-methylamide groups, correspondingly.

The structure was then oriented with OPM^29^ for packing in a membrane and inserted into a POPC bilayer and solvated using Packmol Memgen 1.1.0.^30, 31^ To neutralize the net charge, sodium and chloride ions were added at the concentration of 150 mM. The final molecular system contained nearly 10^5^ atoms and had an approximate volume of 100 × 100 × 100 Å.

The Amber force field parameters were applied to the molecular components of the final structure as follows: ff19SB – to proteins, lipid17 – to lipids, gaff2 – to ligands. The four-point OPC water model was used. Following 5000-step minimization done with the Amber package tool pmemd, the system was heated to 310 K and equilibrated in two stages: under the canonical NVT ensemble for 1 ns, first, and under the isothermal-isobaric NPT ensemble at 1 atm for 1 ns, afterwards. Production simulations at 310 K and 1 atm under the NPT ensemble were then initiated and run for at least 100 ns. Covalent bonds involving hydrogen atoms were constrained using SHAKE. Short-range non-bonded interactions were cut off at 9 Å. The Langevin thermostat and the Berendsen (equilibration) and the Monte-Carlo (production runs) barostats were used. Long-range electrostatic interactions were computed using the Particle Mesh Ewald method.

Accelerated molecular dynamics modeling was performed under the same conditions according to the algorithm described in the Amber manual. Said algorithm speeds up conformational transitions by reducing the energy barriers between the states of the simulated system. Energy threshold and boosting factor parameters were calculated with the formulas presented in the manual.

Publication-quality images were generated in PyMOL.^32^

## 3. Results

### 3.1. Effects of LY294 and LY303 on Ca^2+^ signaling in HEK/5-HT_2C_ cells

Consistently with negligible expression of serotonin receptors,^33^ HEK293 cells were irresponsive to serotonin in terms of Ca^2+^ signaling (Supplemental Fig. 1A). In contrast, HEK/5-HT_2C_ cells responded to nanomolar serotonin with Ca^2+^ transients (please see Supplemental methods 1 and Supplemental Fig. 1B–D for details of the HEK/5-HT_2C_ cells generation and characterization). These serotonin-induced Ca^2+^ responses were reversibly inhibited by LY294 at appropriate doses (Fig. 1A). We generated a LY294 inhibition curve for cells stimulated with 4 nM serotonin and fitted one with Eq. 1 (Fig. 1B), thereby evaluating IC_50_ of the LY294 effects to be 1.2 μM. Being nearly 10-fold lower than LY294 concentrations commonly used for inhibition of PI3K in cell-based assays,^7, 34–36^ this value raised the question whether PI3K was really involved in impairing cell responsiveness (Fig. 1A). Surprisingly, applications of LY303 that is the PI3K-inactive analogue of LY294, triggered agonist-like Ca^2+^ transients with EC_50_ of 2.5 μM (Fig. 1A, C), this unexpected effect of the LY303 could not be mediated by PI3K inhibition. Note that the reported activity of LY294 and LY303 towards PI3K was verified using genetically encoded PIP_3_ reporter PH(Akt)-Venus (Supplemental methods 2 and Supplemental Fig. 2).

**Fig. 1.**
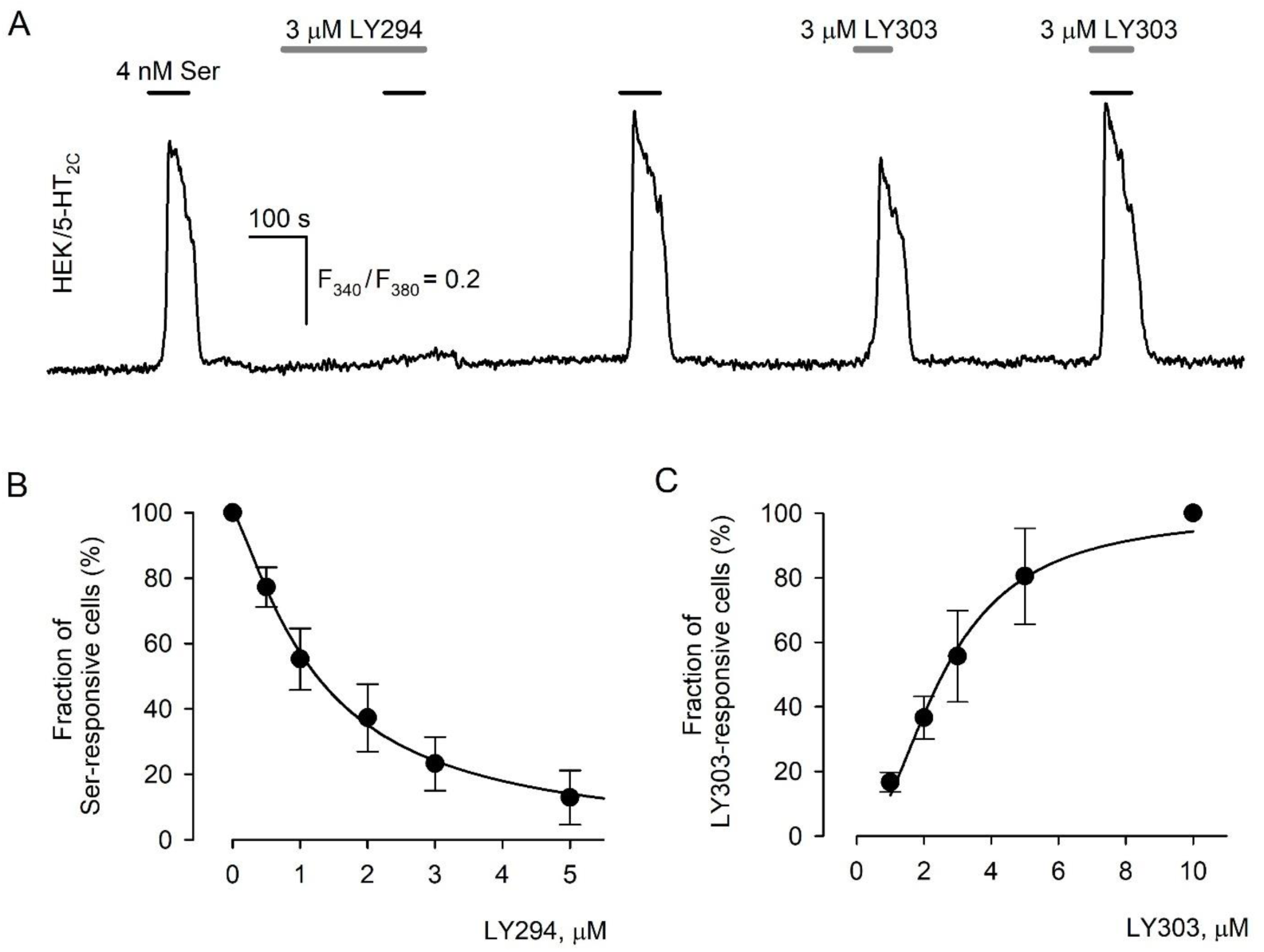
Effects of LY294 and LY303 on Ca^2+^ signaling in HEK/5-HT_2C_ cells. (A) Representative recording of intracellular Ca^2+^ in an individual Fura-2-loaded HEK/5-HT_2C_ cell showing that 3 μM LY294 reversibly inhibited Ca^2+^ signals elicited by 4 nM serotonin, while 3 μM LY303 induced agonist-like Ca^2+^ transients (N = 215). Here and in the below figures, applications of compounds are indicated by the straight-line segments above the experimental traces. Deviations of cytosolic Ca^2+^ in individual cells loaded with Fura-2 are presented as the ratio *F*_340_/*F*_380_, where *F*_340_ and *F*_380_ are the instant intensity of cell fluorescence upon excitation at 340 nm and 380 nm, respectively. (B, C) Fractions of HEK/5-HT_2C_ cells responsive to (B) 4 nM serotonin as a function of LY294 concentration (N = 108–379) and (C) LY303 at varied doses (N = 31–196). The experimental data (filled circles) are presented as a mean ± S.D. The solid curves represent the approximation of the experimental data with the Eq. 1 at *n* = 1.30 and *C_0.5_* = 1.24 (B) and with Eq. 2 at *n* = 2.20 and *C_0.5_* = 2.51 (C).

### 3.2. The effect of LY294 is unrelated to its main known targets PI3K and CK2

To disclose a mechanism responsible for the inhibitory effect of LY294 on serotonin-induced Ca^2+^ transients, we first examined whether it was related to the inhibition of PI3K or casein kinase 2 (CK2), the primary specified targets that are inhibited by LY294 with equal efficacy.^37, 38^ As a control, we used the PI3K and CK2 inhibitors wortmannin and hematein, respectively, which substantially differ from LY294 in chemical structures (Supplemental Fig. 3). It turned out that contrary to LY294, neither wortmannin nor hematein affected serotonin-induced Ca^2+^ transients (Fig. 2A, B). These findings highly devalued the idea that LY294 could affect serotonin responses via inhibition of PI3K and/or CK2 activity, favoring the possibility that some unknown LY294 target was involved.

**Fig. 2.**
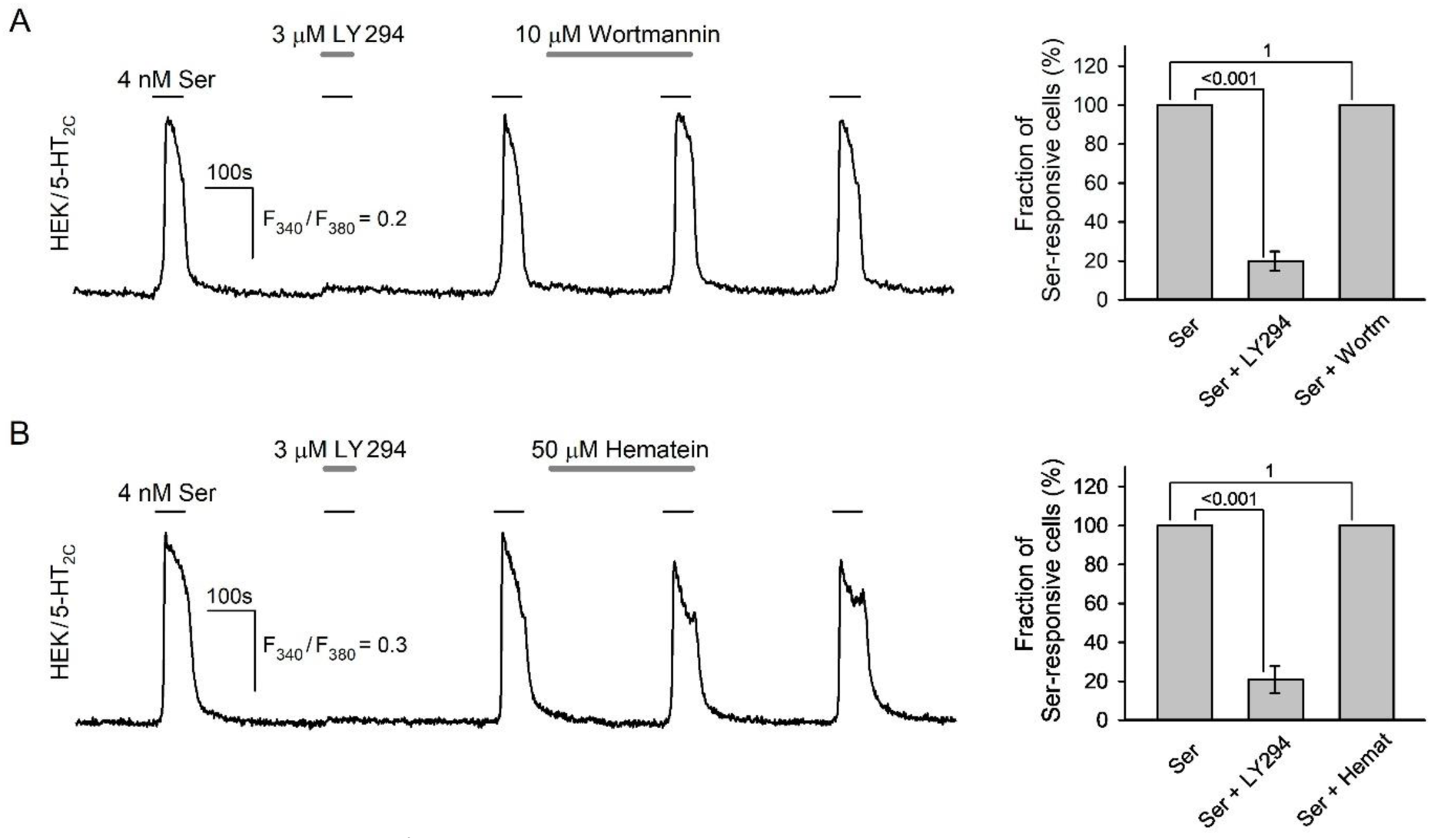
Serotonin-induced Ca^2+^ signals in the presence of LY294, wortmannin, and hematein. (A–B) Representative recordings of serotonin-induced Ca^2+^ transients in individual HEK/5-HT_2C_ cells loaded with Fura-2 *(left panels)* and fraction of cells responsive to serotonin *(right panels)* in control and in the presence of LY294, and (A) wortmannin (N = 150) or (B) hematein (N = 150). Unlike 3 μM LY294, 10 μM wortmannin and 50 μM hematein insignificantly affected cellular responses to 4 nM serotonin. The data in the right panels are presented as a mean ± S.D., one-way repeated measures ANOVA with Holm-Sidak post-hoc test (p values in the graph).

### 3.3. Serotonin 5-HT_2C_ receptor as a target of LY294

Next, we took into account that the LY294 inhibition of cell responsiveness developed for seconds, given that the inhibitor suppressed Ca^2+^ responses even being applied concurrently with serotonin (Fig. 2). This rather rapid effect suggested that LY294 could interfere with transduction machinery directly. If so, a set of putative cellular targets of LY294 would include the 5-HT_2C_ receptor, Gq-protein, PLC, and IP_3_ receptors (IP_3_Rs).

Inquiring first whether LY294 could inhibit IP_3_Rs, we used IP_3_ uncaging to initiate IP_3_-driven Ca^2+^ release directly, that is, not involving the other elements of the phosphoinositide cascade. In designated experiments, the HEK293 cells were loaded with both the Ca^2+^ dye Fluo-8 and the photoconvertible IP_3_ precursor caged-Ins(145)P3. The stimulation of loaded cells with UV flashes photolyzed caged-Ins(145)P3 and produced a jump in cytosolic IP_3_, thereby activating IP_3_Rs and releasing stored Ca^2+^. In control, 32.6 ± 6.1% of the assayed cells generated agonist response-like Ca^2+^ transients upon IP_3_ uncaging, while in the presence of 25 μM LY294, the fraction of these cells increased to 43 ± 8% (Fig. 3A). Thus, in contrast to Ca^2+^ responses to serotonin (Fig. 1A, B), Ca^2+^ release through IP_3_Rs initiated by IP_3_ uncaging was even sensitized by the treatment of cells with LY294. This phenomenon could not be associated with the PI3K inhibition, since LY303 exerted the similar effect (Supplemental Fig. 4A). These findings indicated clearly that IP_3_Rs were not involved in mediating the inhibitory effect of LY294 on responsiveness of HEK/5-HT_2C_ cells to serotonin.

**Fig. 3.**
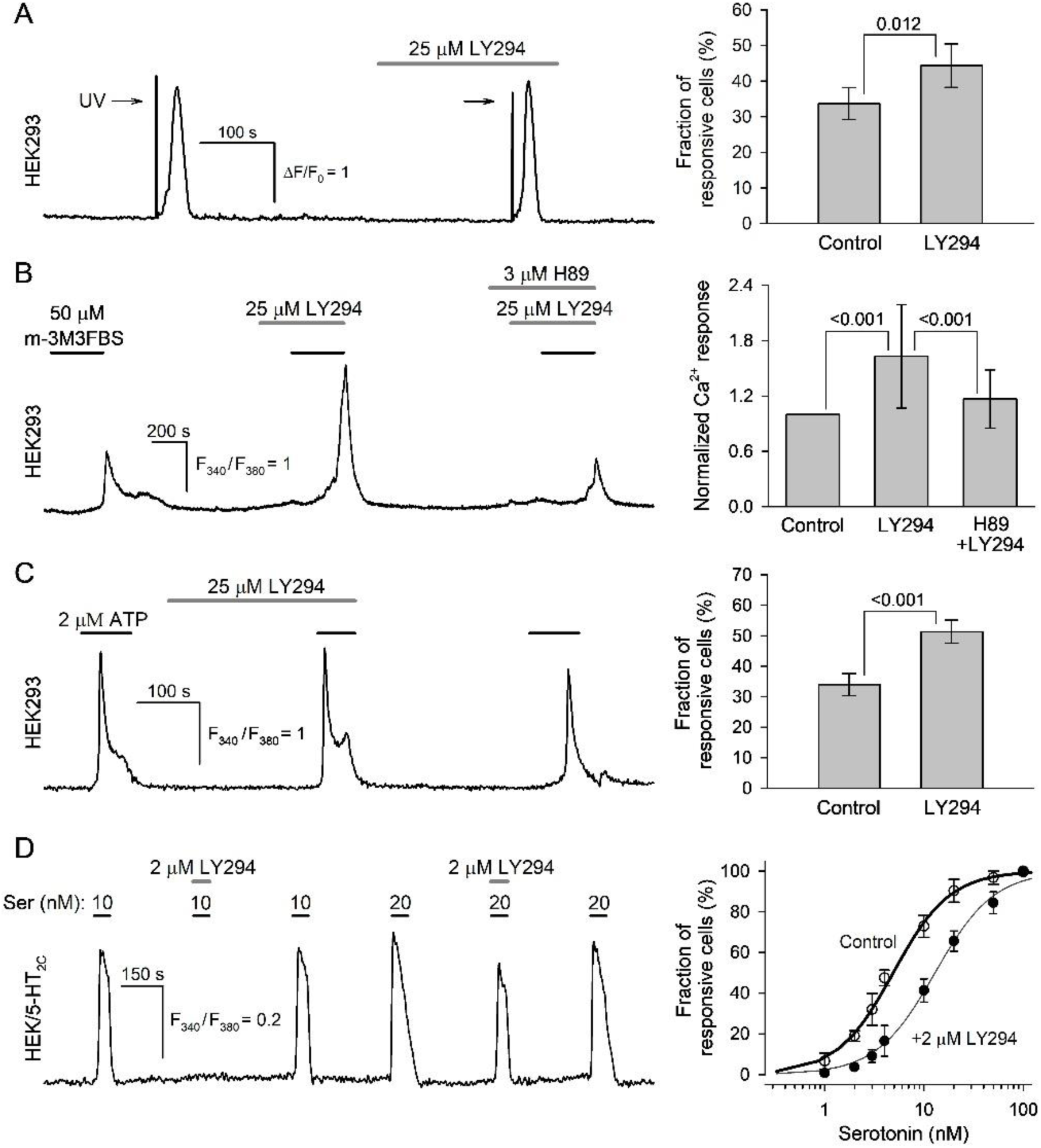
The effects of LY294 on activity of signaling molecules pivotal in serotonin transduction. (A) The effect of LY294 on IP_3_Rs activity. Left panel, representative recording of Ca^2+^ transients elicited by IP_3_ uncaging in an individual HEK293 cell in control and after pretreatment with 25 μM LY294. Сells were loaded with both caged-Ins(145)P3 for IP_3_ uncaging and Fluo-8 for Ca^2+^ imaging. IP_3_ uncaging was stimulated by 0.8-s UV flashes, which produced optical artifacts that appeared as a marked overshoot in the fluorescence traces. Right panel, averaged fractions of cells responsive to IP_3_ uncaging in control and in the presence of 25 μM LY294 (N = 300). Here and in the other right panels, the data are presented as a mean ± S.D. at the indicated numbers of cells. Paired t-test (p values in the graph). (B) The effect of LY294 on PLC activity. Left panel, representative recording of Ca^2+^ signals in an individual HEK293 cell induced by 50 μM PLC activator m-3M3FBS in control and after pretreatment with 25 μM LY294 alone or in combination with 3 μM PKA inhibitor H89. Right panel, normalized magnitudes of Ca^2+^ signals induced by 50 μM m-3M3FBS under the indicated conditions (N = 95). For each individual cell, its response to 50 μM m-3M3FBS in control was taken as a unit. One-way repeated measures ANOVA with Holm-Sidak post-hoc test (p values in the graph). (C) The effect of LY294 on the phosphoinositide cascade triggered by purinergic GPCR. Left panel, representative recording of Ca^2+^ transients induced by 2 μM ATP in an individual HEK293 cell in control and after pretreatment with 25 μM LY294. Right panel, the averaged fractions of ATP-responsive cells (N = 300). Paired t-test (p values in the graph). (D) LY294 as a competitive antagonist of the 5-HT_2C_ receptor. Left panel, representative recording of intracellular Ca^2+^ in an individual HEK/5-HT_2C_ cell. As illustrated, 2 μM LY294 suppressed Ca^2+^ signals elicited by 10 nM serotonin, but failed to cancel responses to 20 nM serotonin (N=148). Right panel, fractions of responsive HEK/5-HT_2C_ cells versus serotonin dose in control (open circles and thick line) and in presence of 2 μM LY294 (filled circles and thin line) (N=532). The curves represent the approximation of the experimental data with the Eq. 2 at *n* = 1.52 and *C_0.5_* = 4.90 (thick line), and at *n* = 1.46 and *C_0.5_* = 12.91 (thin line). In (B–D), cells were loaded with Fura-2.

Considering PLC as one more potential LY294 target, we used the PLC activator m-3M3FBS that was capable of initiating Ca^2+^ signals in HEK293 cells (Fig. 3B), presumably stimulating sufficient IP_3_ production to push Ca^2+^ release through IP_3_Rs. In the presence of LY294, the magnitude of these signals increased by nearly 50% on average, compared to control (Fig. 3B). This LY294 effect also could not be associated with PI3K inhibition as LY303 similarly enlarged magnitudes of cellular responses to m-3M3FBS (Supplemental Fig. 4B). This increase in the magnitude of Ca^2+^ signals could result from the activation of protein kinase A (PKA) by LY294 or LY303,^39^ which in turn could stimulate IP_3_Rs and potentiate the Ca^2+^ release^40^. Indeed, the potentiation of m-3M3FBS responses by LY294 and LY303 was canceled by the PKA inhibitor H89 (Fig. 3B; Supplemental Fig. 4B). Altogether, these observations demonstrated that LY294 did not impair PLC, which therefore could not be responsible for the inhibitory effects of LY294 on serotonin-induced Ca^2+^ transients (Fig. 1A, B).

In the phosphoinositide cascade, the Gq-protein is a signaling molecule operating upstream of PLC and IP_3_Rs. If Gq was a bona fide target of LY294, this compound should also inhibit Ca^2+^ signaling initiated by an agonist of any GPCR coupled to PLC by Gq. Given that HEK293 cells express P2Y receptors^33^ coupled to Gq,^41^ we used the purinergic agonist ATP in the further analysis. Being stimulated by ATP, HEK293 cells generated robust Ca^2+^ signals that were never canceled by LY294 (Fig. 3C). This observation strongly argued against the possibility that activity of the Gq-protein could be inhibited by LY294.

The abovementioned evidence that LY294 did not suppress activity of IP_3_Rs, PLC, and Gq (Fig. 3A–C) allowed one to exclude these signaling molecules from a set of intracellular targets responsible for the collapse of serotonin transduction occurring in the presence of the PI3K inhibitor (Fig. 1A, B). Thus, in the transduction circuit, 5-HT_2C_-Gq-PLC-IP_3_Rs, solely the serotonin receptor remained as a possible LY294 target.

The direct action on the 5-HT_2C_ receptor suggested that LY294 should have exhibited traits of a 5-HT_2C_ antagonist, particularly, somehow distorting a dose-response curve of serotonin responses. We therefore studied responsiveness of cells to serotonin at varied doses in control and in the presence of 2 μM LY294. The dependence was quantified as a fraction of cells responsive to serotonin at a given dose, and in control it was satisfactorily approximated by Eq.2 at EC_50_=4.9 nM and *n*=1.52. Expectedly, 2 μM LY294 weakened cell responsivity, depending on serotonin concentration (Fig. 3D, left panel) and induced a rightward shift of the dose-response curve, albeit without a change in the maximal fraction of serotonin-responsive cells (Fig. 3D, right panel). The LY294-shifted dependence also was well-fitted with Eq.2 at EC_50_= 12.9 nM and *n*=1.46. It thus appears that phenomenologically, LY294 acted as a competitive antagonist of the 5-HT_2C_ receptor.

### 3.4. Serotonin 5-HT_2C_ receptor as a target of LY303

Among the effects described above, the most intriguing was the ability of LY303 to induce Ca^2+^ transients in the HEK/5-HT_2C_ cells (Fig. 1A, C). Note that the reduction of bath Ca^2+^ from 2 mM to 260 nM weakly or negligibly affected LY303-induced responses, which however completely disappeared in the presence of the PLC inhibitor U73122 (Fig. 4A). It thus appeared that the generation of Ca^2+^ responses to LY303 necessitated activation of the phosphoinositide pathway. At the same time, LY303 never initiated Ca^2+^ signaling in parental HEK293 cells (Fig. 4B), indicating that it was unable to stimulate IP_3_ production directly. This finding and that sets of receptors and signaling proteins in HEK/5-HT_2C_ and HEK293 cells differ in the 5-HT_2C_ receptor only, led us to the conclusion that just this GPCR endowed HEK/5-HT_2C_ cells with the responsivity to LY303. This inference was strongly supported by the observation that HEK/5-HT_2C_ cells never responded to LY303 in the presence of the specific 5-HT_2C_ receptor antagonist RS-102221 (Fig. 4C). Amazingly, LY294, whose antagonism towards the 5-HT_2C_ receptor was highlighted above, also suppressed cell responses to LY303 (Fig. 4D). This suggested that most likely, LY303 and LY294 acted on the same target, the 5-HT_2C_ receptor, pointing out that LY303 could serve as a 5-HT_2C_ agonist.

**Fig. 4.**
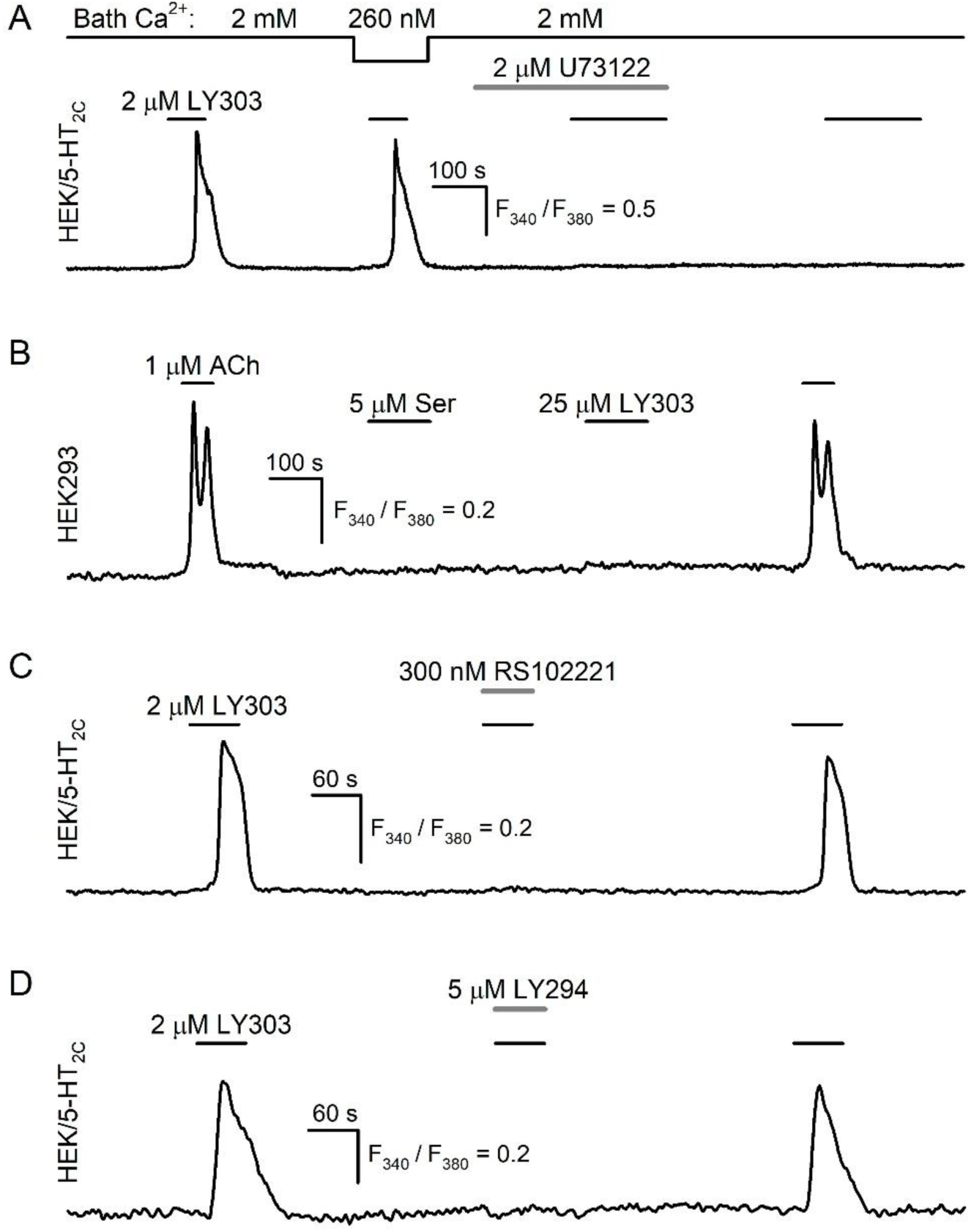
Evidence for LY303 as an agonist of the 5-HT_2C_ receptor. (A–D) Representative recordings of intracellular Ca^2+^ in individual Fura-2-loaded cells. (A) Ca^2+^ signals elicited by 2 μM LY303 in HEK/5-HT_2C_ cells were weakly affected by the reduction of bath Ca^2+^ from 2 mM to 260 nM but those were suppressed by 2 μM PLC inhibitor U73122 (N = 64). (B) Parental HEK293 cells were not responsive to serotonin as well as LY303 with Ca^2+^ transients, while their functionality was validated by robust responses to 1 μM acetylcholine (ACh) (N = 241). (C) Ca^2+^ signals initiated by 2 μM LY303 in HEK/5-HT_2C_ cells were inhibited in the presence of 300 nM 5-HT_2C_ receptor antagonist RS102221 (N = 21). (D) Ca^2+^ signals initiated by 2 μM LY303 in HEK/5-HT_2C_ cells were inhibited in the presence of 5 μM LY294 (N = 36).

### 3.5. Computer modeling of interactions between LY294 and LY303 and the serotonin 5-HT_2C_ receptor

Altogether, our experimental findings (Fig. 1 – Fig. 4) provided indirect evidence that LY294 and LY303 could act as ligands of the 5-HT_2C_ receptor, suggesting the existence of appropriate complexes. To get independent support for this inference, we employed methods of computational biophysics, specifically docking and MD, to examine complexes of the mouse 5-HT_2C_ receptor with LY294 and LY303. Given that the 3D structure of the mouse 5-HT_2C_ receptor has not been resolved yet, we used the structure of the human 5-HT_2C_ receptor as a template (see Materials and Methods). For each ligand, docking was used to localize sites on the surface of the 5-HT_2C_ receptor in which its binding would be most optimal energetically. The stability of docking-predicted ligand-receptor complexes was then examined with MD.

Since a structure of the 5-HT_2C_ receptor in complex with serotonin is currently not available, we relied on the experimentally derived structures of the 5-HT_2C_ receptor in complex with agonist ergotamine (PDB ID: 6BQG) and inverse agonist ritanserin (PDB ID: 6BQH). These structures^20^ suggest that there is a binding site in the receptor transmembrane cavity (Supplemental Fig. 5). To ensure that it was explored as a binding site of LY294 and LY303, the constraining region for docking simulations was chosen as the space bounded by transmembrane helices, extracellular loops, and the geometric center of the receptor.

Docking of serotonin showed that it binds to the 5-HT_2C_ receptor in the same transmembrane cavity (Fig. 5A) as the agonist ergotamine and inverse agonist ritanserin. This docking-predicted location was considered hereafter as an approximation for the 5-HT_2C_ orthosteric site. The evolution of the docking-predicted serotonin-5-HT_2C_ complex was then examined using molecular dynamics. Overall, seven 100-ns MD-trajectories were generated, and all of them showed that the serotonin-5-HT_2C_ complex was quite stable for 100 ns at least (Supplemental Fig. 6). Given, however, that these conventional MD (cMD) trajectories might not be sufficiently long for the complex to reach a relatively rare conformation promoting serotonin release, we also employed the accelerated MD (aMD) approach. By lowering the energy barriers between the states of the simulated system, this algorithm speeds up conformational transitions thus allowing to generate effectively much longer trajectories for the same compute-time. Note that in previous computational studies of 58-residue BPTI protein, the speed-up factor was evaluated to be roughly 10^3^.^42^ However, even aMD-trajectories showed insignificant deviations of serotonin from the initial pose, demonstrating its inability to leave the orthosteric site (Supplemental Fig. 6).

**Fig. 5.**
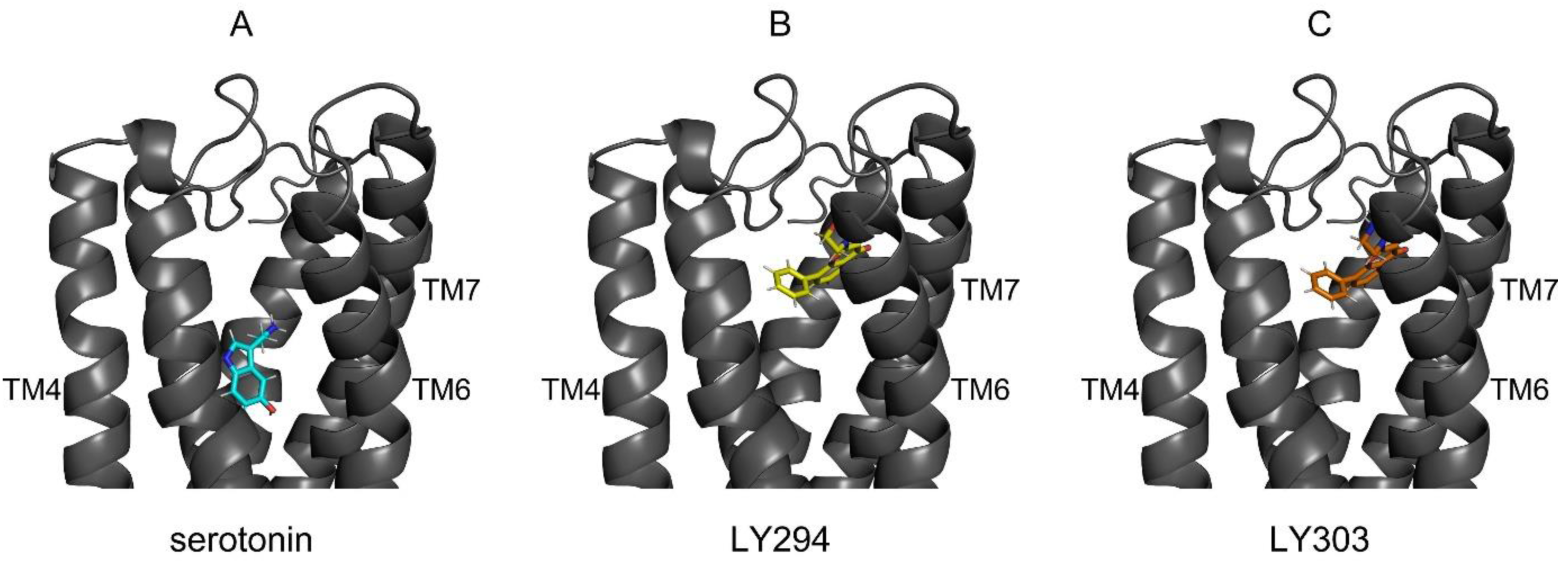
Docking-predicted conformations of the 5-HT_2C_ receptor in complex with serotonin (A), LY294 (B), and LY303 (C). TM4, TM6, and TM7 are 4th, 6th, and 7th transmembrane helices, respectively. As can be seen, both serotonin and the LY-compounds localize in the transmembrane cavity, although serotonin’s position is closer to the geometric center of the receptor than positions of LY294 and LY303. Binding poses of LY294 and LY303 practically coincide, which perhaps is not unexpected, given how structurally similar they are.

Next, we analyzed the interactions of the 5-HT_2C_ receptor with LY294 and LY303. Docking simulations showed that both LY-compounds were similarly bound to the receptor in its orthosteric site (Fig. 5B, C). Both cMD and aMD simulations started with the docking-predicted conformations show the stability of the generated complexes for 100 ns at least (Supplemental Fig. 6). These results support the capability of the LY-compounds to bind to the 5-HT_2C_ receptor, which is necessary for their pharmacological activity, that is, for LY294 to prevent receptor activation by serotonin (Fig. 1) and for LY303 to activate the receptor (Fig. 4).

## 4. Discussion

The PI3K inhibitors wortmannin and LY294 were widely and are still used in cell studies to demonstrate a role for PI3Ks in various intracellular processes. Meanwhile, apart from PI3K, other intracellular targets for LY294 have also been identified. Reportedly, LY294 can affect activity of CK2,^37^ mammalian target of rapamycin (mTOR), ^43^ DNA-dependent protein kinase,^44^ Pim-1 kinase,^45^ nuclear factor-κB (NF-κB),^46^ MRP1,^47^ cAMP phosphodiesterase 2 and 4,^48,49^ slow voltage-gated K^+^ current,^50^ and some other signaling/regulatory proteins.^38^

In a number of studies describing LY294 effects on physiological processes, specific targets remained non-identified. For instance, this PI3K inhibitor elevated cytosolic Ca^2+^ in bovine and human airway smooth muscle (ASM) cells by mobilizing intracellular Ca^2+^ stores, and inhibited subsequent Ca^2+^ responses to carbachol and histamine.^7^ In rat ASM cells, LY294, but not wortmannin, suppressed serotonin-induced Ca^2+^ signals.^8^ LY294 suppressed adenosine-induced release of acetylcholine in the frog neuromuscular junction independently of its effects on PI3K or CK2.^51^ Previously, we also revealed that LY294 and its structural analogs exert the inhibitory effects on Ca^2+^ signaling induced by certain aminergic agonists.^9,10^ In this work we attempted to clarify mechanisms enabling LY294 and its PI3K-inactive analog LY303 to affect Ca^2+^ signaling in HEK/5-HT_2C_ cells expressing the recombinant mouse serotonin 5-HT_2C_ receptor.

The analysis of the inhibitory effect of LY294 on serotonin-induced Ca^2+^ signals provided evidence for a negligible contribution of its specified targets PI3K and CK2 to this phenomenon (Fig. 2). Verifying the idea that LY294 could affect functionality of the serotonin transduction machinery, we sequentially excluded IP_3_Rs, PLC, and Gq, key molecules of the phosphoinositide cascade, as potential LY294 targets (Fig. 3A–C). The exclusion of the phosphoinositide cascade as a whole implied that the 5-HT_2C_ receptor was the most likely target for LY294. Theoretically, LY294 might act as an antagonist of the 5-HT_2C_ receptor or disrupt its coupling to the Gq protein. The first possibility was strongly supported by the fact that LY294 altered the dose-dependence of serotonin in a manner characteristic of a competitive antagonist (Fig. 3D). Consistently, molecular docking and MD simulations showed that LY294 could bind to the 5-HT_2C_ receptor in its orthosteric site (Fig. 5A, B), as should have been the case with a classic competitive antagonist.

It should be mentioned that LY294 alone was used as a PI3K inhibitor in diverse studies, wherein physiological effects of serotonin were assayed. It has particularly been shown that LY294 inhibited serotonin-induced secretion of the neuropeptide sensorin from *Aplysia* sensory neurons, leading to the conclusion that the PI3K pathway is crucial for controlling the long-term facilitation of the *Aplysia* sensorimotor synapse.^52^ In mouse cortical neurons, serotonin-induced phosphorylation of Akt, a primary downstream effector of PI3K, was inhibited by LY294, implicating the PI3K-Akt pathway in mediating this effect.^53^ It also has been demonstrated that LY294 significantly diminished manifestations of the Tourette syndrome induced by the selective 5-HT_2A_/_2C_ agonist (1-(2,5-dimethoxy-4-iodophenyl)-2-aminopropane (DOI)), possibly, by modulating the PI3K/Akt/NF-κB pathway.^54^ Based on effects of LY294, it was inferred that serotonin can suppress PPARγ expression and stimulate proliferation of pulmonary artery smooth muscle cells via the PI3K/Akt pathway, which is associated with pulmonary hypertension development.^55^ In light of our findings, the above-mentioned effects of LY294 may be a result of direct inhibition of serotonin receptors, rather than suppression of PI3K activity.

LY303 is structurally similar to LY294 but inactive towards PI3K,^19^ and it is commonly used as a control verifying the specificity of LY294 effects and, therefore, involvement of PI3K in intracellular processes.^56, 57^ The most surprising observation in our study was that LY303 stimulated Ca^2+^ signaling in the HEK/5-HT_2C_ cells. Taking into account this finding and that the 5-HT_2C_ antagonist RS102221 and LY294 cancelled LY303-induced signals (Fig. 4C, D), we inferred that LY303 acted as an agonist of the 5-HT_2C_ receptor. In support of this idea and consistently with its pharmacological activity, computer simulations showed that LY303 could truly occupy the receptor orthosteric site (Fig. 5C). Given that LY294 and LY303 have a very similar structure (Supplemental Fig. 3) and presumably bind to the same site at the 5-HT_2C_ receptor (Fig. 5B, C), it is unclear why they exerted opposite physiological effects.

This is a complicated question characteristic of receptor pharmacology in general, as the link between ligand binding and activity of a ligand-receptor complex is not unambiguous. It is important to note that a particular ligand can selectively direct receptor movement to a certain area of the conformation space and not to others. Accordingly, an action of a drug affecting a receptor activity is determined not only by affinity but also efficacy, another key parameter.^58^ While the affinity of ligand-receptor binding determines a number of ligand-receptor complexes available at the moment, the efficacy characterizes their fraction capable of initiating a physiological response. For instance, a competitive antagonist is unable to render a receptor active, despite being bound in the orthosteric site, and therefore, their complex has zero efficacy in terms of cell responsiveness. Unfortunately, neither docking nor MD can a priory predict ligand activity. Nevertheless, our theoretical predictions on the direct binding of LY294 and LY303 to the 5-HT_2C_ receptor are consistent with the experimental findings, thereby strongly supporting the idea that these compounds are 5-HT_2C_ ligands.

Finally note that LY303 is believed to have antineoplastic activity. For instance, it has been reported to enhance the sensitivity of cancer cells to drug-induced apoptosis by stimulating H_2_0_2_ production, resulting in mitogen-activated protein kinase activation and up-regulation of death receptors.^59, 60^ LY303 is also known to inhibit cancer cell proliferation via mTOR-dependent and independent mechanisms.^61^ Interestingly, serotonin receptor signaling plays a significant role in the progression of tumors of many types due to its growth-stimulatory effect or suppression of anti-tumor immunity.^62, 63^ Tumor cells express different subtypes of serotonin receptors, including 5-HT_2C_,^63^ and therefore the agonist-like action of LY303 (Fig. 4) should be taken into account, considering its antitumor activity.

In several works antineoplastic potential of LY303 and LY294 was also considered in connection with leukemia,^64^ pancreatic cancer,^65^ and glioma.^66^ So, our evidence for the direct action of LY294 and LY303 on the serotonin 5-HT_2C_ receptor could be useful not only in studying the mechanisms of serotonin-induced signaling, but for detailing the drug safety profile of these compounds and their derivatives.

## 5. Conclusions

In the present work, we demonstrated that the PI3K inhibitor LY294002 suppressed serotonin-induced Ca^2+^ signaling in the HEK/5-HT_2C_ cells, whereas LY303511, its structural analog inactive against PI3K, was capable of initiating agonist-like Ca^2+^ responses. The evidence presented here suggests that these compounds act on the 5-HT_2C_ serotonin receptor directly: LY294002 presumably acts as its antagonist, while LY303511 stimulates its activity. Therefore, our data imply that LY294002 and LY303511 are better to not be used for demonstrating a role of PI3K in intracellular signaling mediated by serotonin receptors, 5-HT_2C_ at least.

## Financial support

This study was supported by the Russian Science Foundation (grant 25-24-00223).

## Conflict of interest

The authors declare no conflicts of interest.

## Data availability

The authors declare that all the data supporting the findings of this study are contained within the paper

## CRediT authorship contribution statement

Polina D. Kotova: Writing – original draft, Visualization, Supervision, Funding acquisition, Conceptualization. Ekaterina A. Dymova: Visualization, Investigation. Oleg O. Lyamin: Writing – original draft, Visualization, Investigation. Olga A. Rogachevskaja: Investigation. Evgeniya A. Voronova: Writing – original draft. Stanislav S. Kolesnikov: Writing – review & editing.

## Supplementary data

## Supplemental methods 1

### Generation and characterization of the НЕК/5HT_2C_ cell line

The recombinant mouse 5-HT_2C_ receptor was expressed in the HEK293 cells, which normally were not responsive to serotonin in terms of Ca^2+^ signaling (Supplemental Fig. 1A). The cells were routinely cultured in the growth medium (1 ml) containing Dulbecco’s modified Eagle’s medium (DMEM; Servicebio), 10% (vol/vol) fetal bovine serum (HyClone), glutamine (1%) and gentamicin (100 μg/ml) (Gibco) in 12-well culture plates. Cells were grown in a humidified atmosphere of 5% CO_2_ at 37°C.

Before transfection, the cells were seeded in 12-well culture plates at a density of 3–5 ×10^5^ cells per well in 800 μl growth medium and allowed to attach overnight. The transient transfection was performed by adding 100 μl of OPTI-MEM (Gibco) containing 1 μg plasmid DNA (pDsRed-Monomer/5-HT_2C_)^1^ and 2 μl GenJect-39 (Molecta) to cultured cells. After 24-h incubation, the transfection mixture was replaced with the growth culture medium, and 24– 96 h after transfection cells were assayed physiologically to confirm the correct functioning of the 5-HT_2C_ receptor.

To generate a stable cell line, the transfected cells were cultured in the growth medium supplemented with 0.8 mg/ml G-418 (Invivogen) for 3 weeks, resulting in a cell population containing about 70% 5-HT_2C_-positive cells. The obtained stable cell line НЕК/5HT_2C_ was further cultured in the presence of 0.3 mg/ml G-418.

The serotonin responsiveness of the HEK/5-HT_2C_ cells in terms of Ca^2+^ signaling was then tested. Individual HEK/5-HT_2C_ cells were irresponsive to serotonin applied at subnanomolar concentrations but generated Ca^2+^ transients of similar magnitudes at different doses should they exceed the threshold (Supplemental Fig. 1В, C). Thus, serotonin stimulated Ca^2+^ signaling in HEK/5-HT_2C_ cells in an “all-or-nothing manner”, so their sensitivity to it could not be characterized by Ca^2+^ response magnitude. In this study, we commonly used an alternative dose dependence, evaluating a fraction of cells responsive to serotonin at a particular concentration (Supplemental Fig. 1D). The EC_50_ value of serotonin was calculated by fitting this dose dependence using the Eq. 2 (see the main text).

**Supplemental Fig. 1.**
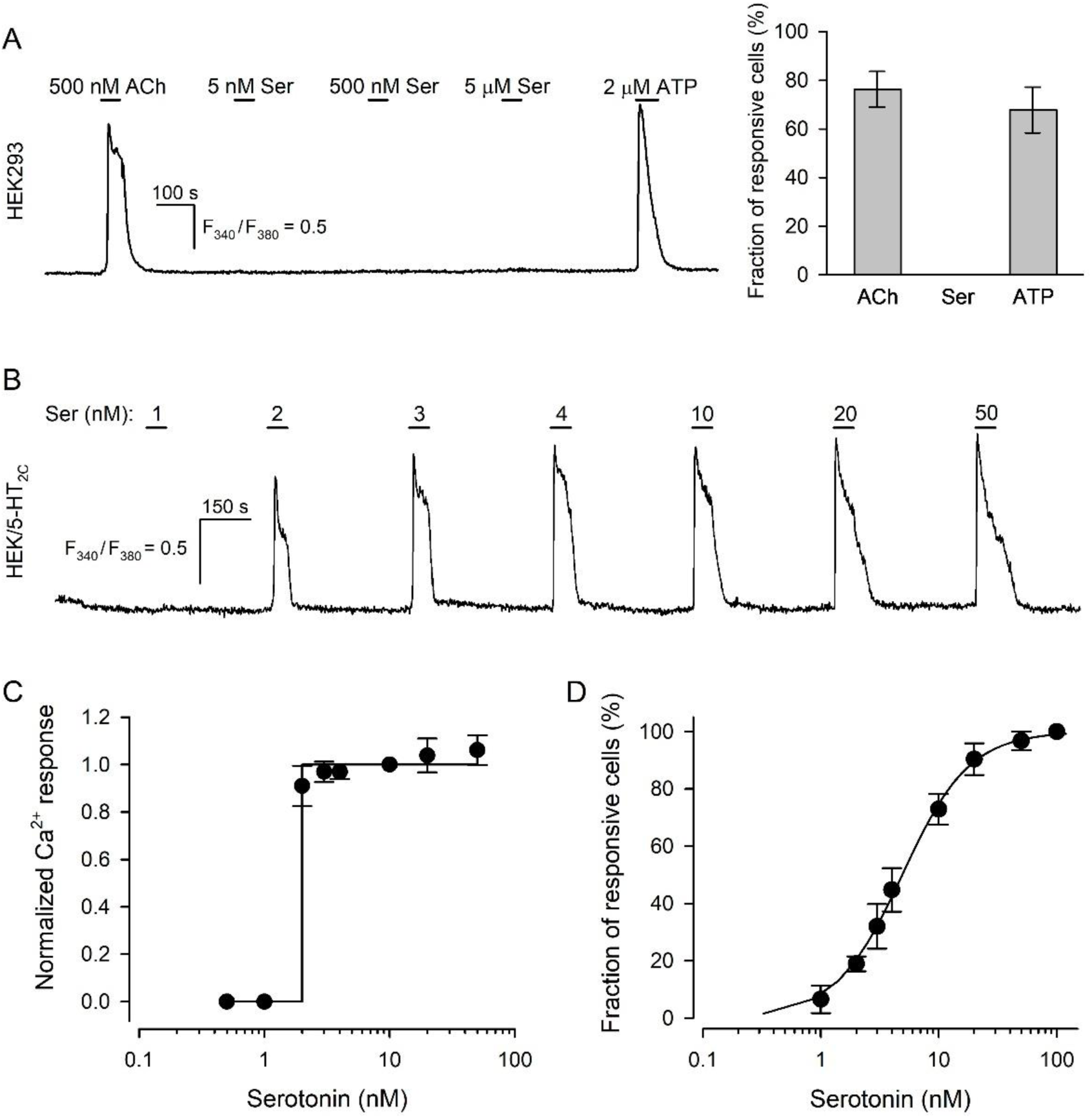
Responsiveness of the HEK293 and HEK/5-HT_2C_ cells to serotonin. (A) Representative recording of Ca^2+^ responses in an individual Fura-2-loaded HEK293 cell to sequentially applied acetylcholine (ACh), serotonin (Ser), and ATP at the indicated doses (*left panel*), and corresponding fractions of responsive HEK293 cells (N = 300) (*right panel*). The data are presented as a mean ± S.D. The number of all tested cells in each experiment was set as 100%. (B) Representative recording of Ca^2+^ responses in an individual Fura-2-loaded HEK/5-HT_2C_ cell sequentially stimulated by serotonin at the indicated doses. (C) Averaged magnitude of Ca^2+^ response versus serotonin dose obtained for HEK/5-HT_2C_ cells exhibiting the same threshold of 2 nM serotonin (N = 10). For a given cell, a response at a particular serotonin concentration was normalized to that evoked by 10 nM serotonin. The experimental data (filled circles) were approximated by the Heaviside function with a jump at 2 nM (solid line). The experimental data are presented as a mean ± S.D. (D) Dose dependence of the fraction of serotonin-responsive HEK/5-HT_2C_ cells (N = 532). The number of cells responsive to 100 nM serotonin was set as 100%. The experimental data (filled circles) are presented as a mean ± S.D. The solid curve represents the approximation of the experimental data with the Eq. 2 at *n* = 1.52 and *C_0.5_* = 4.90.

## Supplemental methods 2

### Verification of the LY294 and LY303 activity towards PI3K

To verify activity of LY294 and LY303 against PI3K, we used monoclonal cell line HEK/R-GECO1/PH(Akt)-Venus, expressing the genetically encoded PIP_3_ reporter PH(Akt)-Venus.^2^ The basis for monitoring PI3K activity with this reporter was as follows. In a resting cell fluorescence of PH(Akt)-Venus is generally distributed among the cytosol almost uniformly. When PI3K is activated and elevates a PIP3 level in the plasmalemma, PH(Akt)-Venus is translocated from the cytosol to plasma membrane, and its redistribution is visible in fluorescent light.^3^

For PIP_3_ imaging, HEK/PH(Akt)-Venus cells were cultured in a hand-made photometric chamber for 24 h prior to experiments in the complete growth medium. The chamber was a Plexiglas framework with an ellipsoidal slot of nearly 150 μl volume, the bottom of the chamber was a disposable coverslip (Menzel-Glaser). Experiments were carried out using an inverted fluorescence microscope Axiovert 200 equipped with an objective Plan NeoFluar 20x/0.75 (Carl Zeiss), a digital sCMOS camera Zyla 4.2P (Andor Technology), metal halide light source AMH-200-F6S (Andor Technology), and spinning disk for confocal microscopy Revolution DSD2 (Andor Technology). Fluorescence of PH(Akt)-Venus was excited at 500 ± 10 nm, emission was collected at 535 ± 15 nm. Serial fluorescence images were captured with NIS-Elements 5.30.01 (Nikon), depending on an experimental protocol. All chemicals were applied by the complete replacement of the bath solution in a 150 μl photometric chamber for nearly 2 s using a perfusion system driven by gravity.

Note that PIP_3_ production in HEK/PH(Akt)-Venus cells could be initiated by insulin via its tyrosine kinase receptor coupled to the PI3K/Akt pathway.^4^ The protocol of the experiment is shown on Supplemental Fig. 2A. Expectedly, PH(Akt)-Venus fluorescence was distributed almost uniformly in non-stimulated cells, excepting for certain narrow zones with slightly increased emission presumably associated with plasma membranes (Supplemental Fig. 2B, left panel). Next, cells were treated with 30 μM LY294 and then stimulated with 100 nM insulin but no marked change in the distribution of PH(Akt)-Venus fluorescence was observed (Supplemental Fig. 2B, middle panel). After removal of LY294 from the bath followed by a rinse of cells, applied once again insulin was at this time capable of initiating the translocation of PH(Akt)-Venus to the plasmalemma. This was evident from the decreased emission of cell bodies and appearance of relatively bright and contrast narrow zones related to plasma membranes of assayed cells (Supplemental Fig. 2B, right panel). In the identical assay, LY303 at the same dose did not prevent insulin-induced translocation of PH(Akt)-Venus to the plasmalemma (Supplemental Fig. 2C). These observations indicated clearly that LY294 inhibited PI3K indeed, whereas LY303 exerted no detectable effect on its activity.

**Supplemental Fig. 2.**
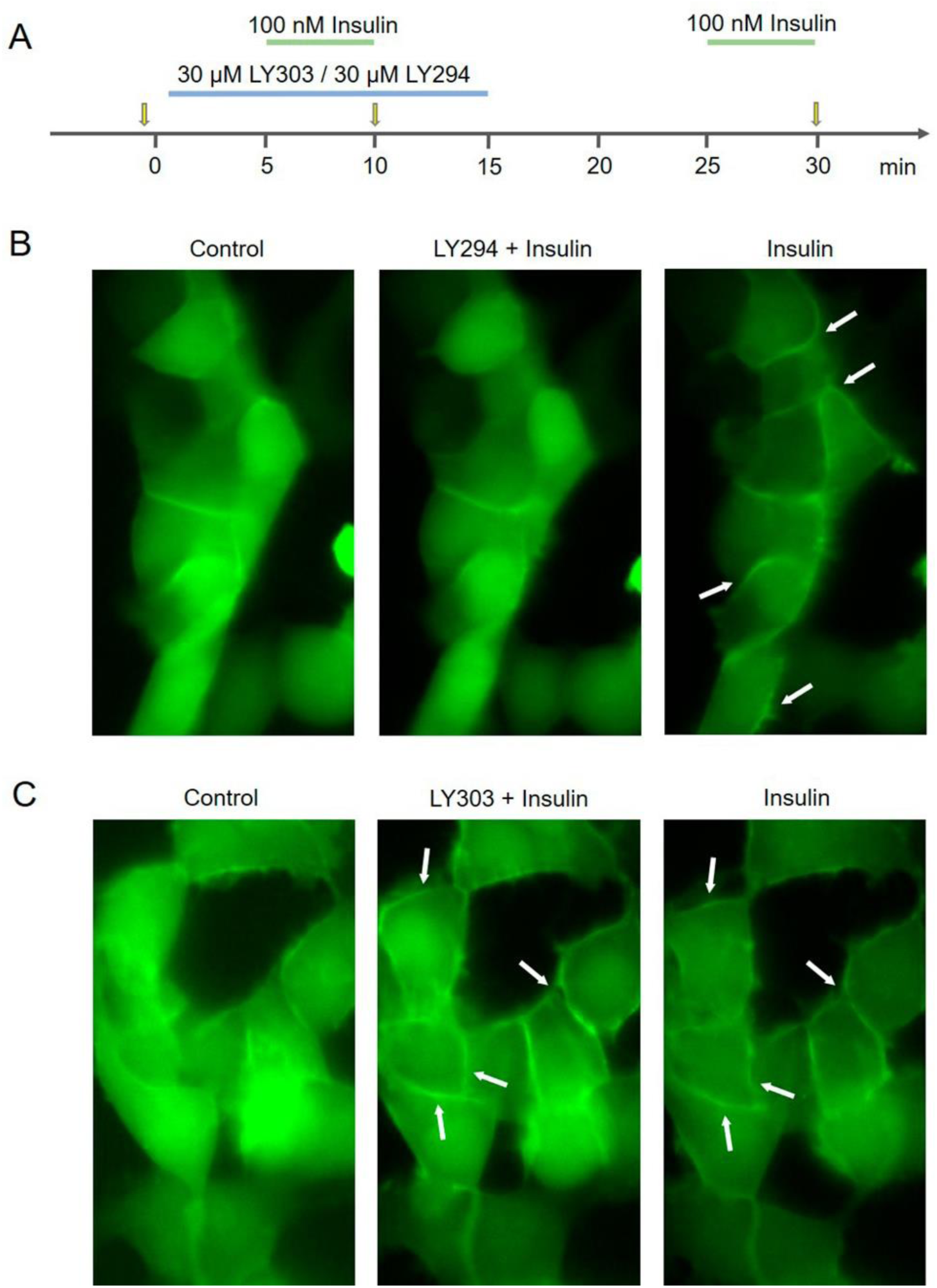
Activity of LY294 and LY303 against PI3K. (A) Experimental protocol timing. The applications of LY294 or LY303, and insulin are indicated by the line segments above the time axis; the moments of image capturing are indicated by the yellow arrows. (B, C) Representative sequential images of HEK/PH(Akt)-Venus cells obtained at the moments indicated in (A). *The left panels*, control images of the group of cells obtained right before drug application demonstrate the virtually homogeneous distribution of PH(Akt)-Venus fluorescence over cell bodies. *The middle panels*, stimulation of cells with 100 nM insulin in the presence of 30 μM LY294 negligibly affected PH(Akt)-Venus distribution compared to control (B), while in the presence of 30 μM LY303 insulin initiated a drop in the fluorescence of the cell cytosol and the appearance of detectable stripe-like fluorescent zones indicated by the white arrows (C), suggesting the insulin-induced accumulation of the PH(Akt)-Venus sensor in the plasmalemma. *The right panels,* images obtained after LY294 or LY303 washing out and subsequent stimulation with insulin.

## Supplemental figures

**Supplemental Fig. 3.**
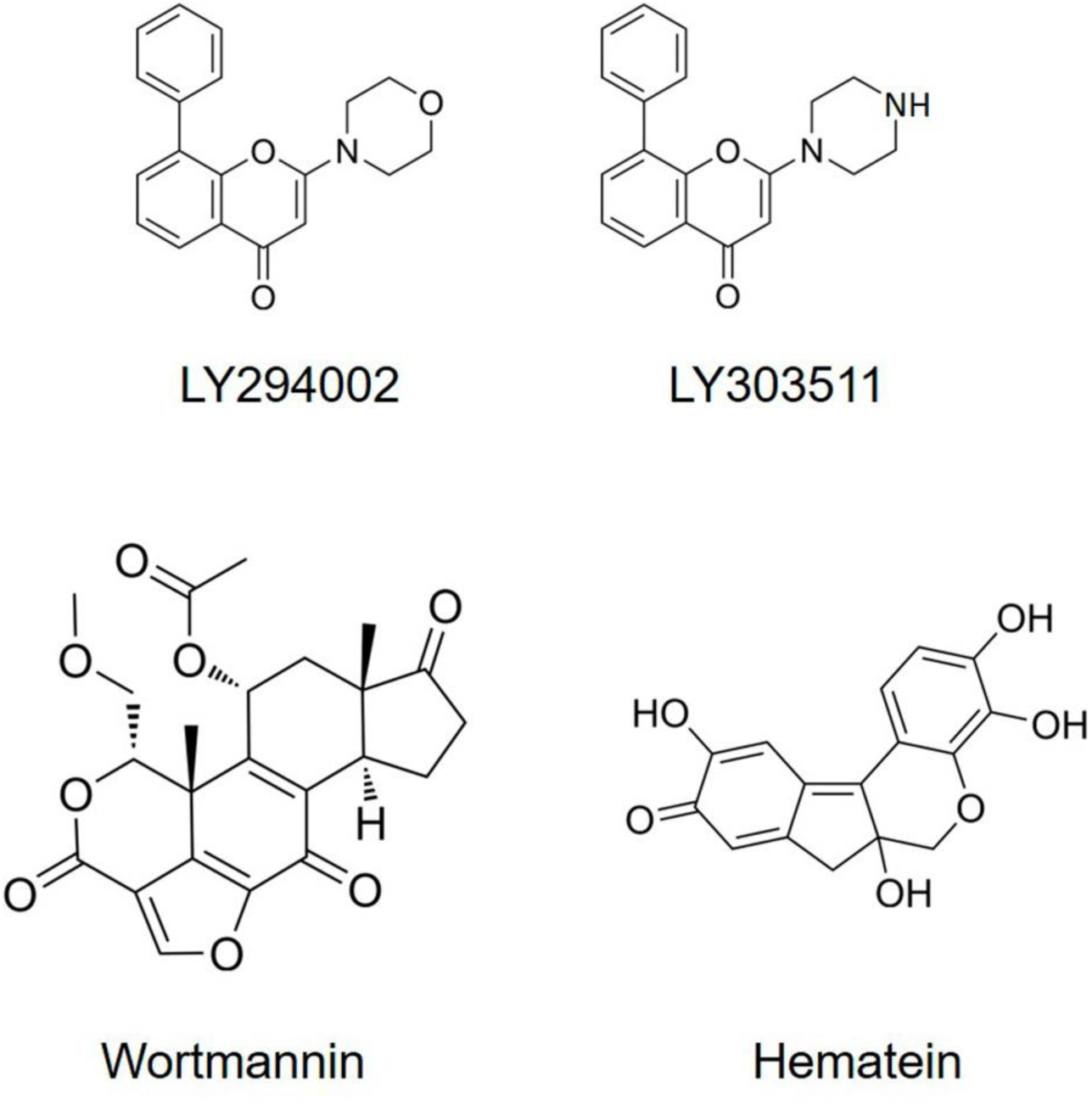
Structural formulas of the PI3K inhibitors wortmannin and LY294002, its PI3K-inactive structural analogue LY303511, and CK2 inhibitor hematein.

**Supplemental Fig. 4.**
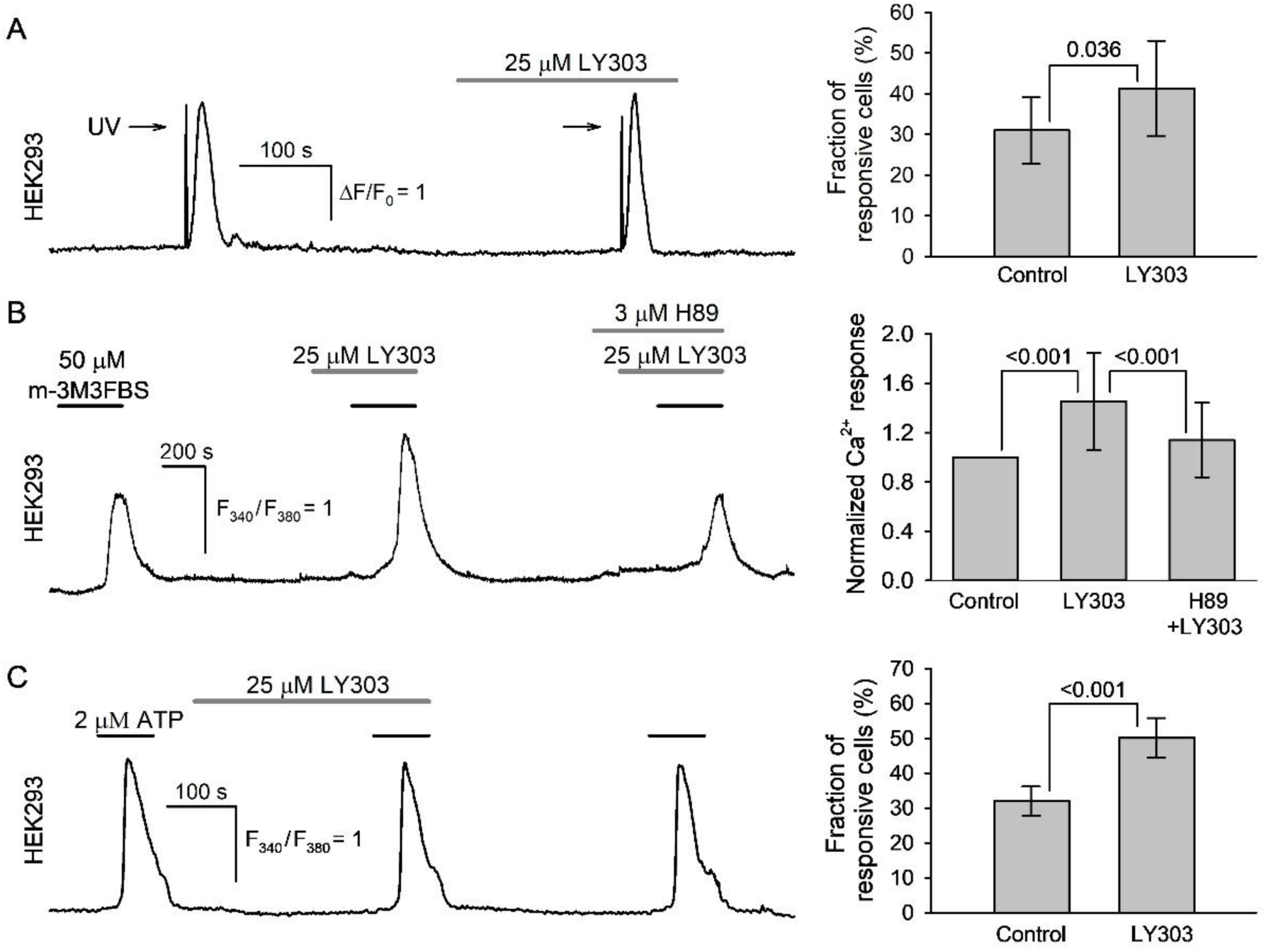
The effects of LY303511 on activity of signaling molecules pivotal in serotonin transduction. (A) The effect of LY303511 (LY303) on IP_3_Rs activity. Left panel, representative recording of Ca^2+^ transients elicited by IP_3_ uncaging in an individual HEK293 cell in control and after pretreatment with 25 μM LY303. Сells were loaded with both caged-Ins(145)P3 for IP_3_ uncaging and Fluo-8 for Ca^2+^ imaging. IP_3_ uncaging was stimulated by 0.8-s UV flashes, which produced optical artifacts that appeared as a marked overshoot in the fluorescence traces. Right panel, averaged fractions of cells responsive to IP_3_ uncaging in control and in the presence of 25 μM LY303 (N = 300). Here and in the other right panels, the data are presented as a mean ± S.D. at the indicated numbers of cells. Paired t-test (p values in the graph). (B) The effect of LY303 on PLC activity. Left panel, representative recording of Ca^2+^ signals in an individual HEK293 cell induced by 50 μM PLC activator m-3M3FBS in control and after pretreatment with 25 μM LY303 alone or in combination with 3 μM PKA inhibitor H89. Right panel, normalized magnitudes of Ca^2+^ signals induced by 50 μM m-3M3FBS under the indicated conditions (N = 65). For each individual cell, its response to 50 μM m-3M3FBS in control was taken as a unit. One-way repeated measures ANOVA with Holm-Sidak post-hoc test (p values in the graph). (C) The effect of LY303 on the phosphoinositide cascade triggered by purinergic GPCR. Left panel, representative recording of Ca^2+^ transients induced by 2 μM ATP in an individual HEK293 cell in control and after pretreatment with 25 μM LY303. Right panel, the averaged fractions of ATP-responsive cells (N = 300). Paired t-test (p values in the graph).

**Supplemental Fig. 5.**
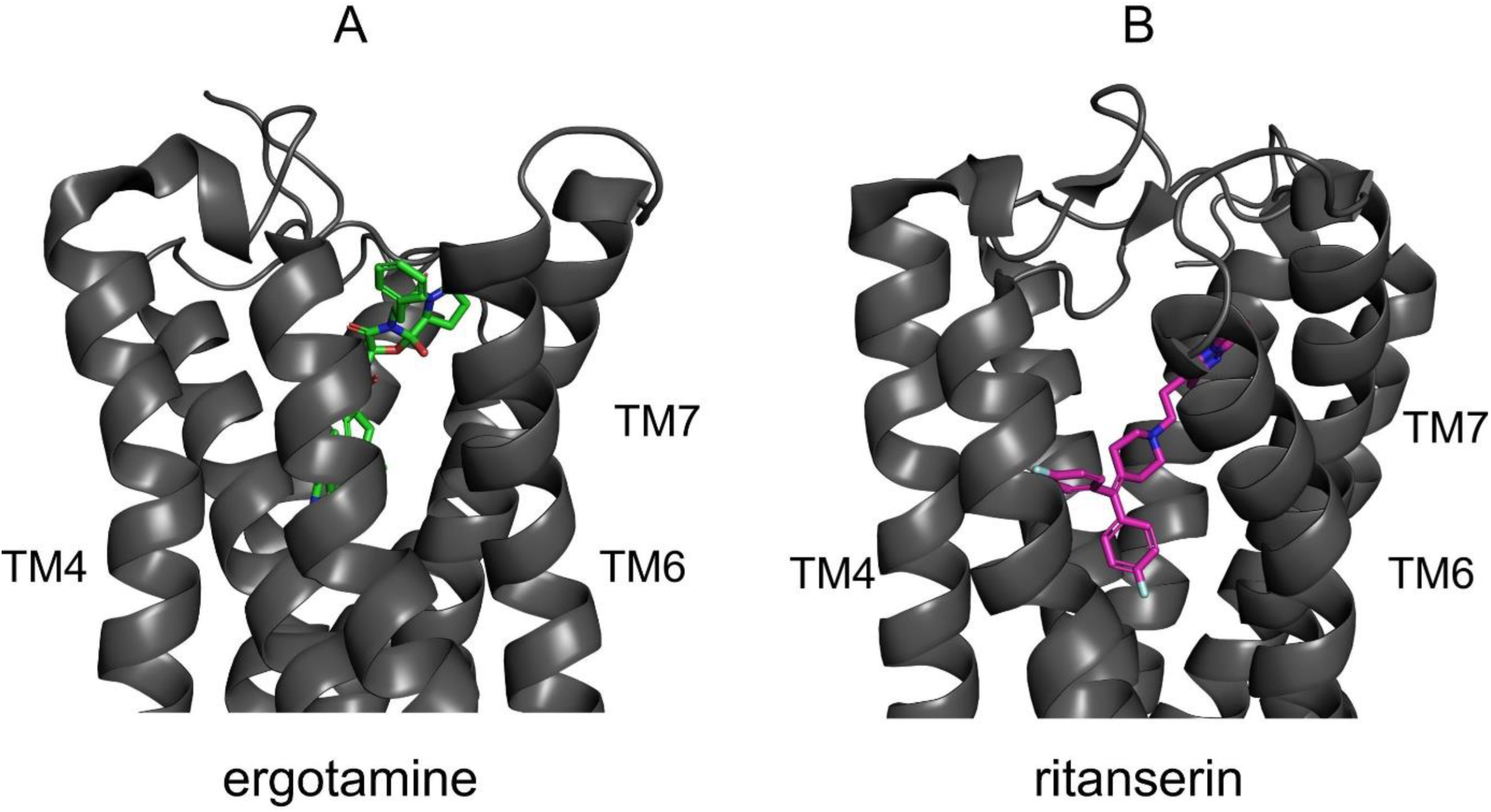
Structures of the 5-HT_2C_ receptor in complexes with the agonist ergotamine and inverse agonist ritanserin derived experimentally.^5^

**Supplemental Fig. 6.**
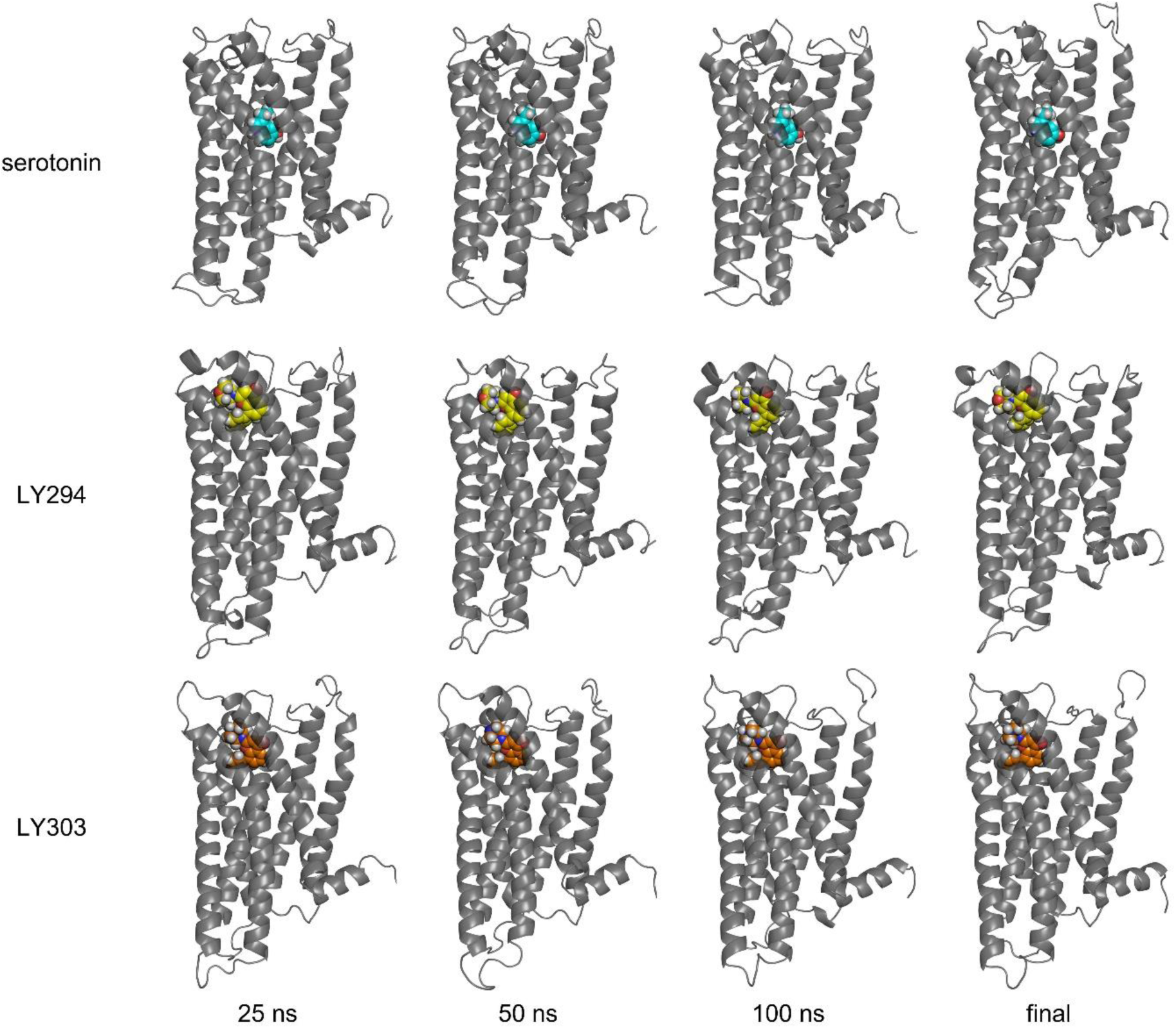
Molecular dynamics snapshots taken from the trajectories at 25, 50, and 100 ns generated for the 5-HT_2C_ receptor in complexes with serotonin, LY294, and LY303. Frame labeled as “final” represents the end state of the structures after modeling with accelerated molecular dynamics. In all simulations, docking-predicted conformations were used as the initial structures As can be seen, serotonin and the LY-compounds by and large remained in their initial locations.

## Notes

### Competing Interest Statement

The authors have declared no competing interest.

